# Genetic estimates of the initial peopling of Polynesian islands actually reflect later inter-island contacts

**DOI:** 10.1101/2022.12.01.518673

**Authors:** Yilei Huang, Shai Carmi, David Reich, Harald Ringbauer

## Abstract

The timing of the initial peopling of the Polynesian islands remains highly debated. Suggested dates are primarily based on archaeological evidence and differ by several hundred years. Ioannidis et al. [2021] used genome-wide data from 430 modern individuals from 21 Pacific islands to obtain genetic estimates. Their results supported late settlement dates, e.g. approximately 1200 CE for Rapa Nui. However, when investigating the underlying model we found that the genetic estimator used by Ioannidis et al. [2021] is biased to be about 300 years too old. Correcting for this bias gives genetic settlement dates that are more recent than any dates consistent with archaeological records, as radiocarbon dating of human-modified artifacts shows settlement definitively earlier than the bias-corrected genetic estimates. These too-recent estimates can only be explained by substantial gene flow between islands after their initial settlements. Therefore, contacts attested by archaeological and linguistic evidence [Kirch, 2021] must have been accompanied also by demographically significant movement of people. This gene flow well after the initial settlements was not modelled by Ioannidis et al. [2021] and challenges their interpretation that carving anthropomorphic stone statues was spread during initial settlements of islands. Instead, the distribution of this cultural practice likely reflects later inter-island exchanges, as suggested earlier [Kirch, 2017].

## Main Text

To estimate settlement times, Ioannidis et al. [2021] analyzed long DNA segments inherited from a common ancestor (identical by descent, henceforth referred to as IBD). After inferring these IBD segments from genome-wide DNA data for pairs of individuals on different Polynesian islands and restricting the analysis to genomic regions of Polynesian ancestry, they fitted the observed IBD length distribution with an exponential curve. From the decay constant, they computed the number of generations that elapsed since the divergence of the two island populations. The bias in the method used by Ioannidis et al. [2021] is that this model assumes that the time to the most recent common ancestor (TMRCA) of all IBD segments shared between samples of any two islands is the same as the split time of their ancestral population. However, the TMRCA is guaranteed to be older than the population split time: at the time of the population split, the ancestors of the two contemporary lineages almost certainly belonged to different people. Therefore, the genetic divergence time also includes the coalescence time of two lineages within the ancestral population before the population split, which as we show in what follows, introduces a substantial overestimate of split times.

To quantify this effect, we derived an analytic expression for the IBD length distribution (Supplement Eq. 4) building on a long-standing theoretical framework [Palamara et al., 2012, Ralph and Coop, 2013, Palamara and Pe’er, 2013, Carmi et al., 2014, Browning and Browning, 2015, Ni et al., 2015, Ringbauer et al., 2017]. We also simulated IBD under this model (1.3). The exact equation and the simulations agree, and both deviate markedly from the model assumed in Ioannidis et al. [2021] (Fig. S1). Importantly, our results show that the methodology employed by Ioannidis et al. [2021] yields a systematic over-estimate of split times of about 10 generations under a wide range of plausible demographic scenarios (Fig. 1), including the ones inferred by Ioannidis et al. [2021]. When assuming a generation time of about 30 years, this bias translates to about 300 years. Correcting the genetic split time estimates of Ioannidis et al. [2021] for this bias gives dates much younger than any archaeologically plausible settlement model.

**Figure 1:**
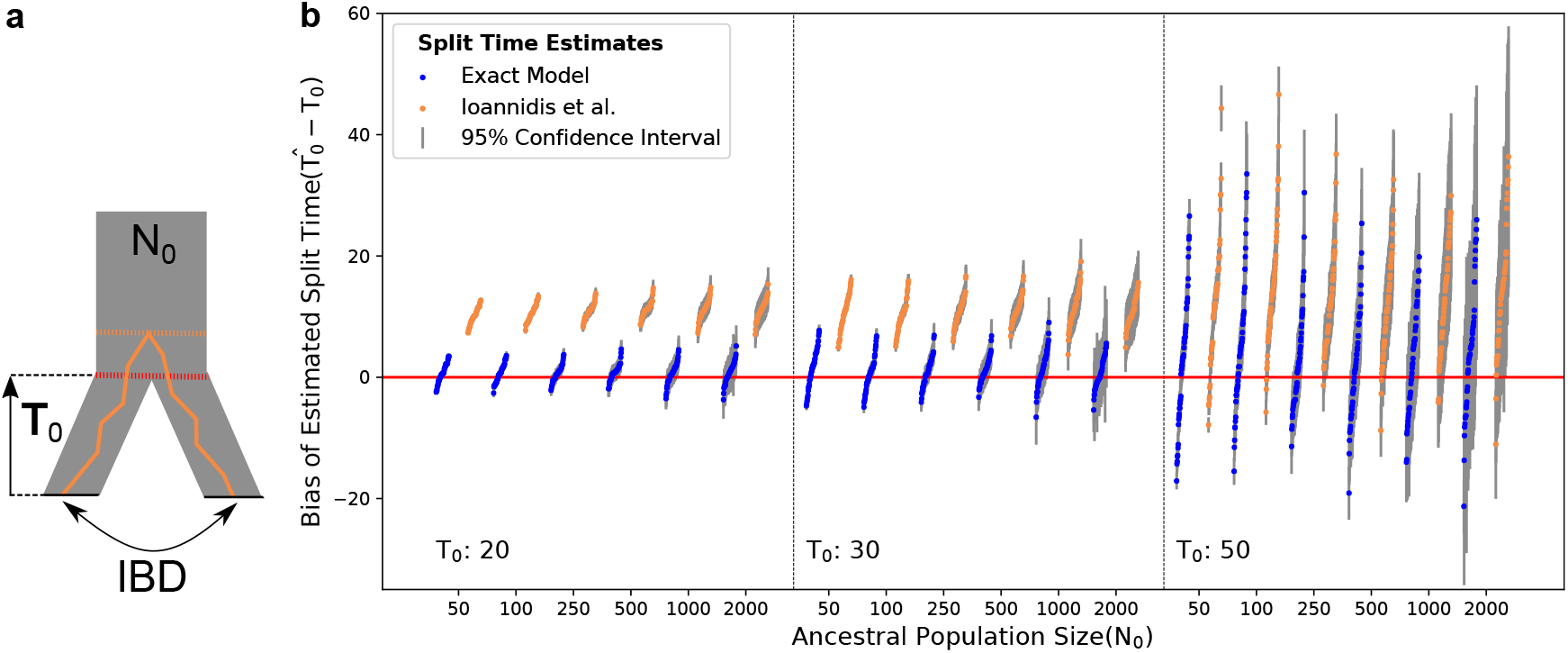
Estimating split times from IBD when simulating Two-Island Models. We fitted both the two-island model and the model in [Ioannidis et al., 2021] to IBD segments simulated under a two-island split model without subsequent gene flow. We simulated all combinations of three different split times *T*_0_ = 20, 30, 50 and six different ancestral population sizes *N*_0_ = 50, 100, 250, 500, 1000, 2000, each with 50 independent replicates recording IBD between two samples of 15 diploid individuals per island. We plotted the biases of the estimated split time 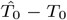 for each of the simulated scenarios.

One process not modelled by Ioannidis et al. [2021] is gene flow postdating the initial settlements of Polynesia. Such contacts introduce shared IBD segments, biasing split time estimates to be too recent. To ameliorate this problem, Ioannidis et al. [2021] restricted their analysis to segments shorter than 15 centimorgan. While this filtering removes long IBD segments that are highly enriched to be of recent origin, recent gene flow introduces IBD segments across all length scales (Fig. S3) and an effect on split time estimates remains (Fig. S4). Major postsettlement cultural exchange almost certainly occurred given linguistic evidence that suggests “considerable and continuing inter-island contact during the early period of eastward migration into the Eastern Polynesian archipelagos” and archaeological evidence of long-distance exchange of stone tools between early Eastern Polynesians [Kirch, 2021]. The genetic split time estimates dating well after initial settlement make it clear that later contacts between islands must have involved substantial movements of people.

Ioannidis et al. [2021] point out that in the case of post-settlement gene flow, their inferred genetic divergence dates between Polynesian islands are in theory only the most recent possible dates of settlement (‘terminus ante quem’). However, they argue that in practice they effectively estimate the dates of settlement for the remote island groups. As support for this claim, they invoke a prior archaeological hypothesis, writing ‘In the case of the most remote islands such as Rapa Nui, which are believed to have had no large-scale population exchanges with other islands, the IBD-based date should coincide closely with the actual date of settlement.’ But here we find that the bias-corrected divergence dates between remote islands actually post-date well-accepted radiocarbon dates of those islands by hundreds of years, including also for Rapa Nui. Therefore, the interpretation of Ioannidis et al. [2021] that IBD-based split times reflect settlement dates on remote islands cannot be correct. Instead, the IBD-based genetic divergence dates must be dominated by previously underappreciated major gene flow events well after initial settlement, including for remote islands such as Rapa Nui.

This evidence of substantial gene flow well after initial settlement based on combining genetic and archaeological evidence brings into question one of the most remarkable implications of Ioannidis et al. [2021] regarding the origins of carving monumental anthropomorphic stone statues, a cultural practice shared among the far-flung islands of Marquesas, Raivavae, and Rapa Nui. It was previously argued that those and other exemplars of stone working reflect inter-island contacts well after the initial settlements [Kirch, 2017]. Ioannidis et al. [2021] conjectured instead that these practices spread into all these islands with their first inhabitants, plausibly from a source in the Tuamotu Islands. The fact that genetic clustering coincides with the practice of carving of monumental statues in Polynesia is an important observation, but our results imply that this signal was likely misinterpreted by Ioannidis et al. [2021], and instead was driven by later and equally extraordinary inter-island exchanges.

## Code Availability

The code used for simulating IBD and analyzing simulated IBD data is publicly available at https://github.com/hyl317/two_island.

## Acknowledgments

We thank Alexander Ioannidis, Patrick Kirch, Johannes Krause, Andres Moreno-Estrada, Kathrin Nägele, and Mark Stoneking for helpful comments on drafts of this manuscript.

## Author Contributions

We annotate author contributions using the CRediT Taxonomy labels (https://casrai.org/credit/).

- Conceptualization (Design of study) – YH, HR
- Software – YH
- Formal Analysis – YH
- Writing (original draft preparation) – YH, DR, SC, HR
- Writing (review and editing) – YH, DR, SC, HR
- Supervision – HR

## 1 Supplementary Information

### 1.1 The exact analytical Model for IBD sharing

Here, we derive an exact expression for IBD sharing in a two-island model with a split with no subsequent gene flow (see model in Fig. 1a). We use an established theoretical framework to exactly model IBD sharing [Palamara et al., 2012, Ralph and Coop, 2013, Palamara and Pe’er, 2013, Carmi et al., 2014, Browning and Browning, 2015, Ni et al., 2015, Ringbauer et al., 2017]. Using the notation of Ringbauer et al. [2017], the expected number of IBD segments 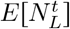 of length *L* and with TMRCA *t*, can be expressed as:

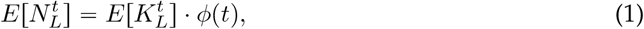

where 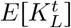 denotes the expected number of overlapping ancestry blocks *L* Morgan long at time *t* before sampling, and *ϕ*(*t*) denotes the single-locus coalescent time distribution, i.e. the probability that two lineages coalesce at time *t* ago. Throughout, the notation *E*[·] denotes a probability density with respect to time or/and IBD segment length [Ringbauer et al., 2017]. To obtain an analytical expression for the two-island split model, we give expression for both factors in Eq. 1.

First, as described in [Ringbauer et al., 2017], modelling recombination as a Poisson process along a chromosome of length *G* yields:

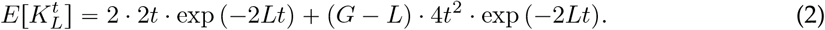

Second, we plug in the coalescent time distribution *ϕ* (*t*) for the specific demographic model of a two-island split with no subsequent gene flow (Fig. S1a). Denoting the split time as *T*_0_ generations back and the ancestral population size as *N*_0_ yields:

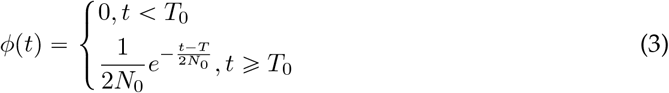

To calculate *E*[*N*_*L*_], the density of expected number of IBD segments of length *L*, we integrate 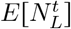 over *t*. To simplify notation, let *β* = 1 + 4*N*_0_*L*. Plugging in Eq. 2 yields the full result:

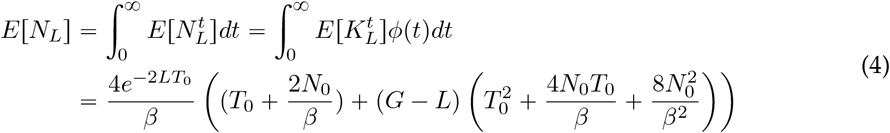

We note that for *G* ≫ *L*, this expression can be approximated by the second term only (effectively ignoring IBD on the chromosome edges). Here, we used the exact full expression throughout, effectively including chromosomal edge effects, as there is little computational overhead.

### 1.2 Inferring demographic parameters by fitting IBD

To estimate demographic parameters from IBD blocks, we apply the method described in Appendix C of Ringbauer et al. [2017]. Summarizing briefly, the expected number of IBD segments within a small length bin [*L, L* + *δ*_*L*_] is given by *δ*_*L*_ · *E* [*N*_*L*_]. Within each length bin, we assume the number of segments to be Poisson distributed around mean *δ*_*L*_ *E* [*N*_*L*_]. We then approximate the total likelihood to be the product of Poisson likelihoods over all length bins, assuming that each length bin is independent. To jointly estimate *T*_0_ and *N*_0_, we maximize this likelihood by using the L-BFGS-B subroutine [Zhu et al., 1997] provided in SciPy[Virtanen et al., 2020]. We then estimated the standard error of the MLE estimate of split time and ancestral population size by numerically calculating the Fisher Information of the likelihood function using the Python package numdifftools, and then approximated the 95% confidence interval by ±1.96 × standard errors.

### 1.3 Simulating IBD sharing

To validate formula Eq. 4, we compared predicted IBD sharing to simulated IBD sharing. To simulated a IBD dataset, we used the coalescent simulator msprime [Kelleher et al., 2016] for chromosome 5 using the HapMap genetic map[Consortium et al., 2005]. We simulated all combinations of *T*_0_ = 20, 30, 50 and *N*_0_ = 50, 100, 250, 500, 1000, 2000. For each demographic model we drew one haplotype from each of the two islands, and simulated 50,000 independent replicates. To keep runtime manageable, we terminated the simulation at 1000 generations as coalescent events before that are extremely unlikely to result in IBD segments several centimorgan long.

We recorded IBD segments according to the full ancestral recombination graph (ARG). Specifically, we define an IBD segment to be a contiguous segment where the most recent common ancestor(MRCA) remains the same without any recombination node on the branch leading up to coalescence. We note that some recombination events do not change the pairwise TMRCA, but our procedure records all IBD segments including those truncated by these ineffective recombinations that would not be detectable in practice. Doing so is usually a good approximation: While ineffective recombination leads to conflation of short IBD segments, especially in populations with very small effective sizes [Chiang et al., 2016], for most realistic demographic parameters and longer IBD blocks these effects remain negligible [Chiang et al., 2016, Ringbauer et al., 2017].

For simulated IBD segments 5-15 cM long, the full theoretical prediction (Eq. 4) matches the simulated data (Fig.S1). Moreover, the simulated IBD distribution exhibits a substantial positive curvature when using a log-scale for the y axis, demonstrating that the decay deviates from a simple exponential model (which would be a straight line on a log scale).

### 1.4 Inferring Split Times from simulated IBD

Next, we evaluated IBD-based demographic inference on simulated data. As described above, we used msprime to simulate data on all 22 autosomes for all combinations of *T*_0_ = 20, 30, 50 and *N*_0_ = 50, 100, 250, 500, 1000, 2000. For each demographic model, we performed 50 independent simulations. In each replicate simulation, we drew 15 diploid samples from each of the two islands and recorded their IBD segments according to the simulated full ARGs. Similar as above, we defined IBD segments to be a contiguous genomic segment along which the MRCA stays constant without any recombination events.

To infer split time and ancestral population size, we used segments 5-15cM long and set the bin size to 0.1*cM*. We calculated the likelihood based on the exact formula for expected IBD sharing (Eq. 4). We noticed that likelihood optimization occasionally converges to only a local optimum when far from the true values, so we started the optimization procedure from three different initial values ((25, 100), (25, 2000), (25, 10000) where the first element is the initial search value for *T*_0_ and the second element is the initial search value for *N*_0_) and returned the output that gives the highest likelihood. For comparison, we also estimated split times using the MLE formula for truncated exponential distribution, following the procedure described in [Ioannidis et al., 2021].

Our results indicate that fitting an exponential decay as in [Ioannidis et al., 2021] results in a systematic over-estimate of split time by about 10 generations across a wide range of demographic parameters (Fig.1). In contrast, fitting the full formula generally yields estimates of both split time and ancestral population sizes with little biases (Fig. 1). We note that for both models, the variance of the estimate increases with larger *T*_0_ or *N*_0_, as do the estimated confidence intervals. This behavior is not surprising, because larger *T*_0_ or *N*_0_ results in fewer shared IBD blocks and inference based on less data tends to be more variable.

## Supplementary Figures

**Figure S1:**
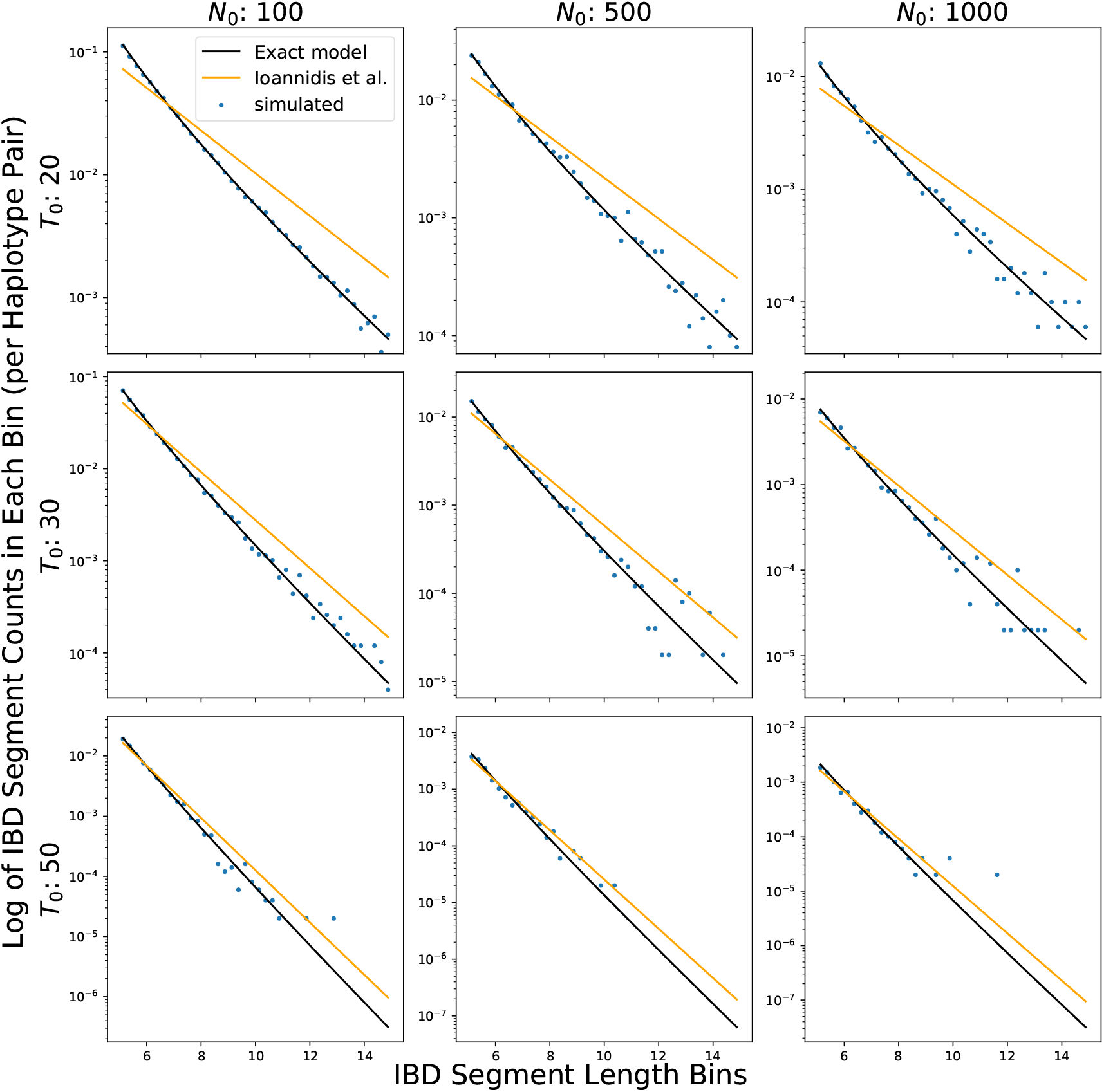
IBD segment distribution in the Two-Island Split Model. We simulated two-island split model with various split times *T*_0_ (rows) and various population sizes *N*_0_ (columns). We show the amount of IBD segments of different lengths (blue dots) between two islands. The exponential fit line (yellow) is calculated as in Ioannidis et al. [2021], we calculated the normalizing constant for the exponential decay curve by matching the total number of observed IBD segments.

**Figure S2:**
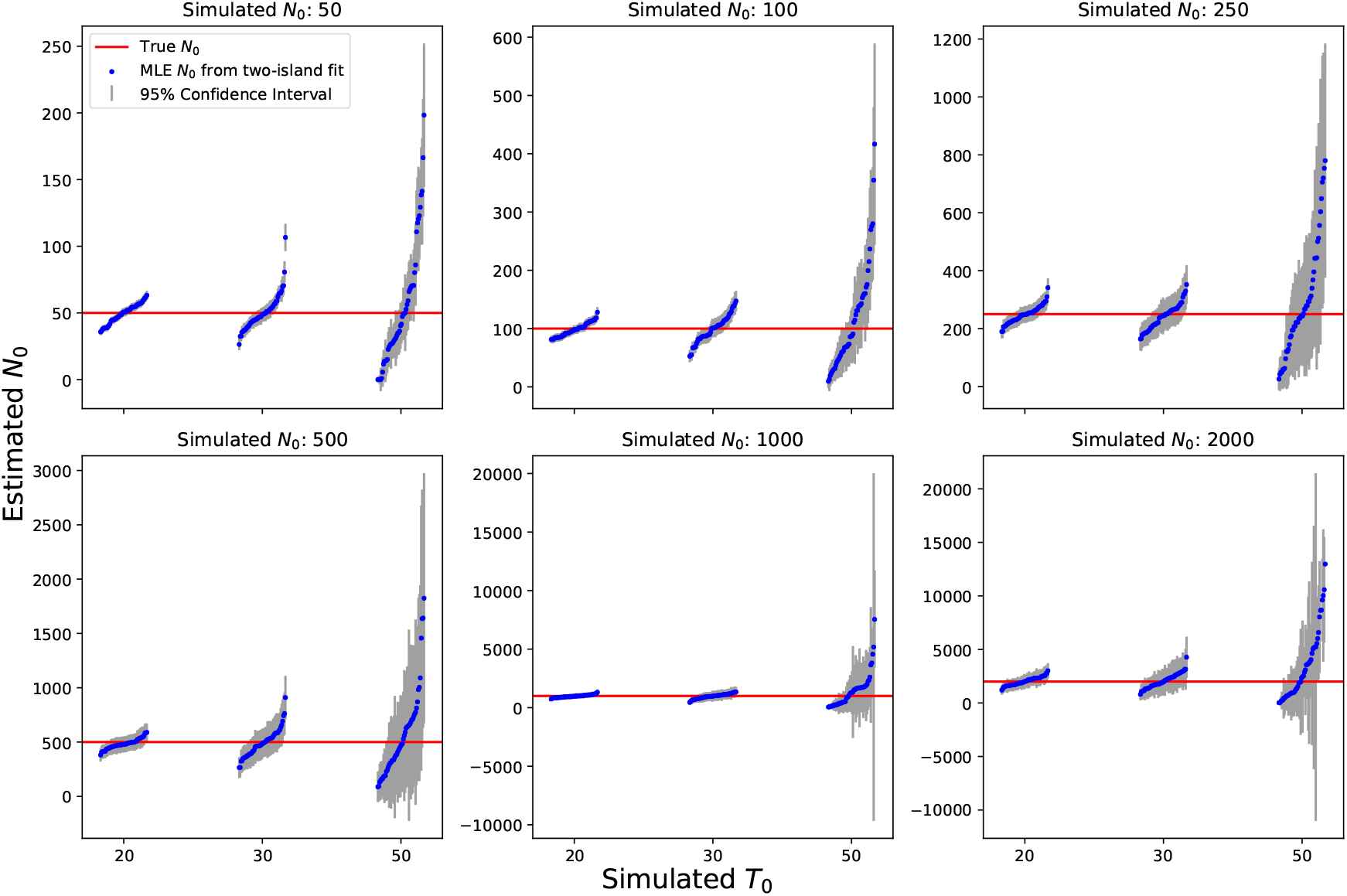
Estimating ancestral effective Population Sizes for Two-Island Split Models. We applied our two-island model to IBD segments simulated for two-island split models to estimate ancestral population sizes before the split. Each panel shows the estimates of one simulated population size (blue dots) with different simulated split times across panels.

**Figure S3:**
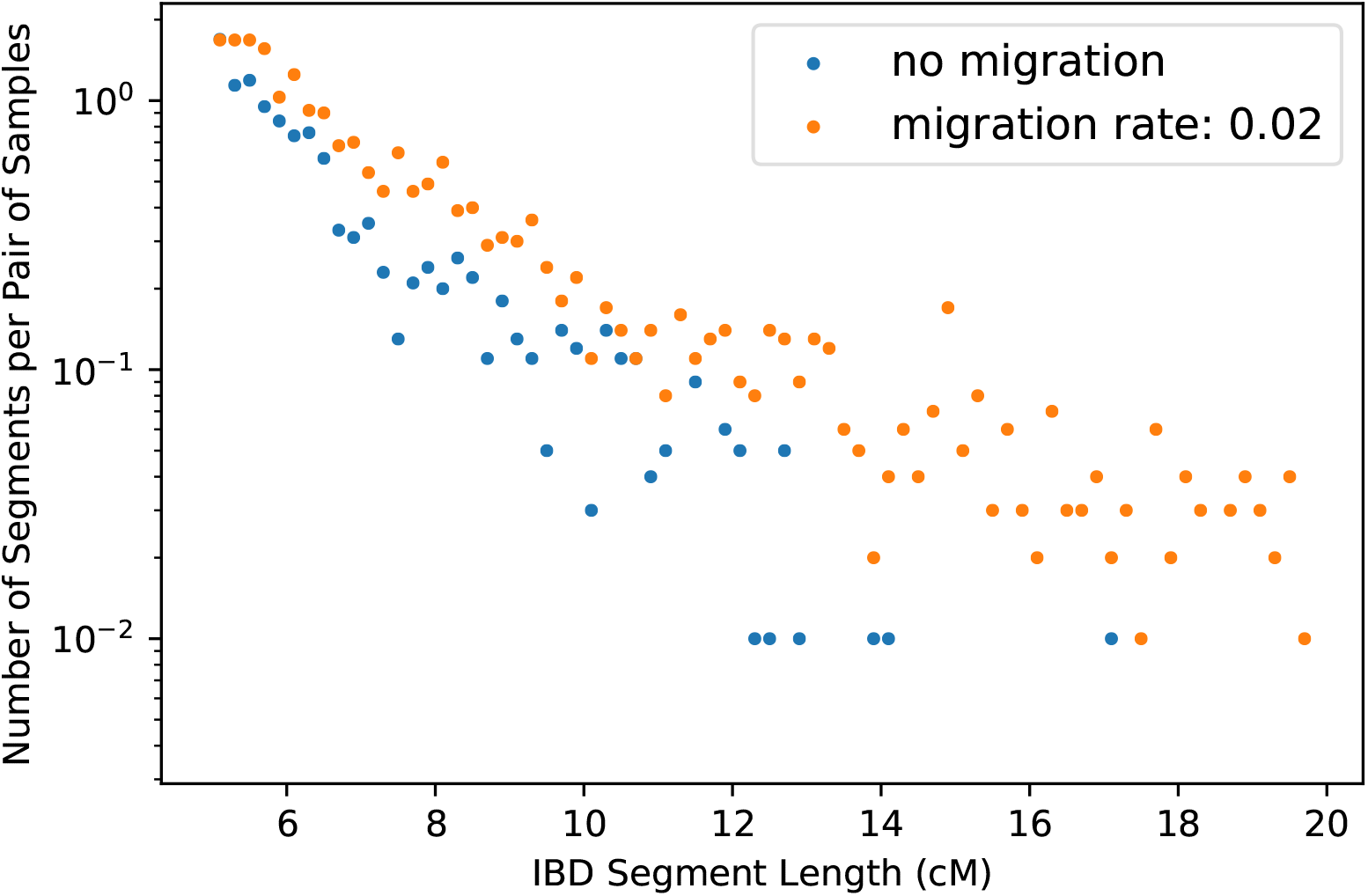
Comparing IBD Length Distributions with and without Migration. Histogram of IBD sharing simulated under the two-island split model (*T* = 20, *N* = 500), with no migration (m=0, blue) and a 2% migration rate (m=0.02, orange) per generation after the split.

**Figure S4:**
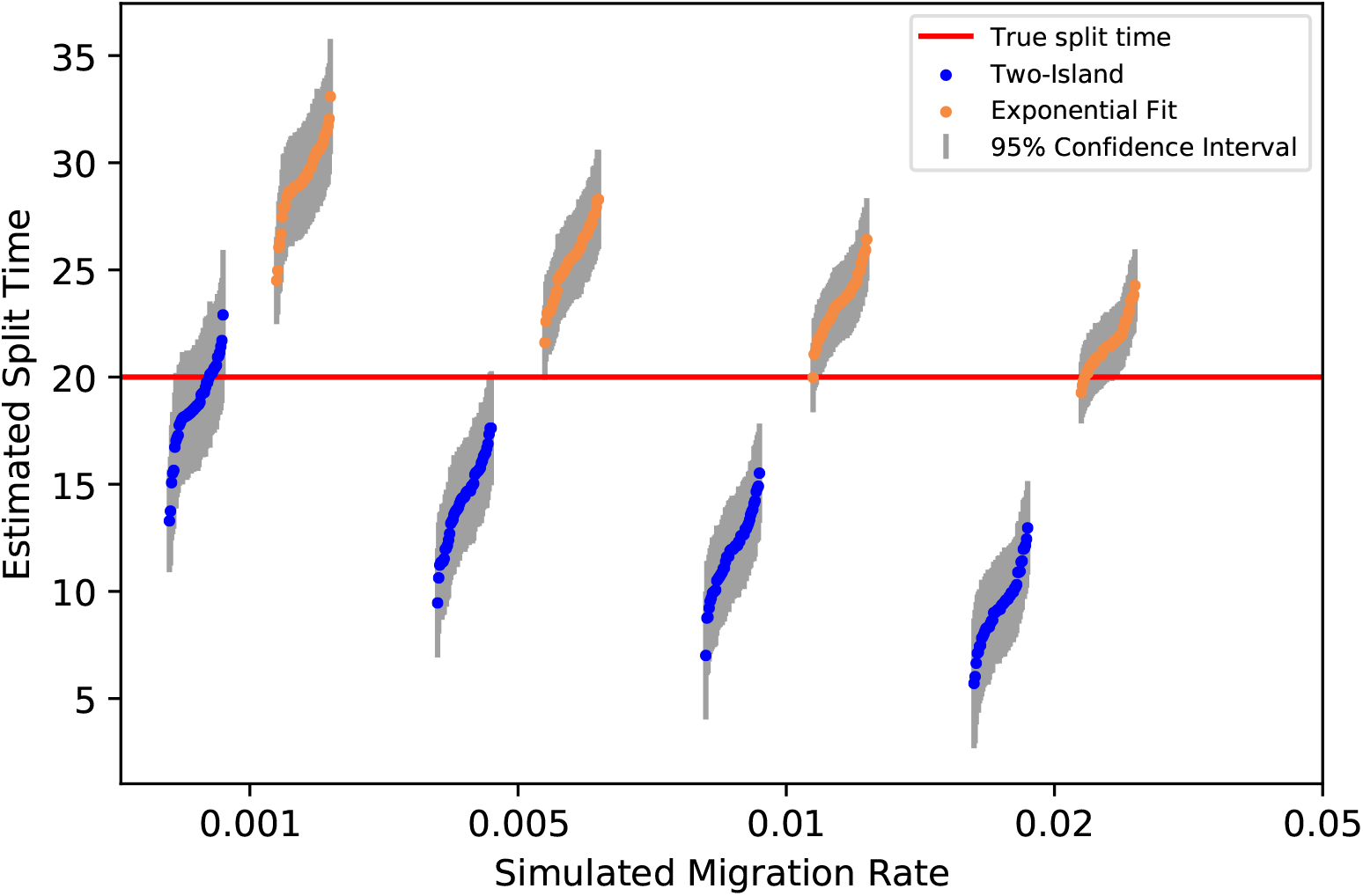
Estimating Split Times with Simulated Gene Flow when assuming none. We simulated a two-island split model with split time *T*_0_ = 20 and ancestral population size *N*_0_ = 1000 as described in the main article, except that we added varying levels of gene flow (m=0.001-0.02) and fit the estimated split times using the exponential model (orange) and the exact two-island model assuming a clean split (blue).

## References

Sharon R Browning and Brian L Browning. Accurate non-parametric estimation of recent effective population size from segments of identity by descent. The American Journal of Human Genetics, 97(3):404–418, 2015.

Shai Carmi, Peter R Wilton, John Wakeley, and Itsik Pe’er. A renewal theory approach to ibd sharing. Theoretical Population Biology, 97:35–48, 2014.

Charleston WK Chiang, Peter Ralph, and John Novembre. Conflation of short identity-by-descent segments bias their inferred length distribution. G3: Genes, Genomes, Genetics, 6(5): 1287–1296, 2016.

International HapMap Consortium et al. A haplotype map of the human genome. Nature, 437 (7063):1299, 2005.

Alexander G Ioannidis, Javier Blanco-Portillo, Karla Sandoval, Erika Hagelberg, Carmina Barberena-Jonas, Adrian VS Hill, Juan Esteban Rodríguez-Rodríguez, Keolu Fox, Kathryn Robson, Sonia Haoa-Cardinali, et al. Paths and timings of the peopling of Polynesia inferred from genomic networks. Nature, 597(7877):522–526, 2021.

Jerome Kelleher, Alison M Etheridge, and Gilean McVean. Efficient coalescent simulation and genealogical analysis for large sample sizes. PLoS computational biology, 12(5):e1004842, 2016.

Patrick V Kirch. Modern polynesian genomes offer clues to early eastward migrations, 2021. Patrick Vinton Kirch. On the road of the winds: an archaeological history of the Pacific Islands before European contact. Univ of California Press, 2017.

Xumin Ni, Wei Guo, Kai Yuan, Xiong Yang, Zhiming Ma, Shuhua Xu, and Shihua Zhang. A probabilistic method for estimating the sharing of identity by descent for populations with migration. IEEE/ACM Transactions on Computational Biology and Bioinformatics, 13(2):281–290, 2015.

Pier Francesco Palamara and Itsik Pe’er. Inference of historical migration rates via haplotype sharing. Bioinformatics, 29(13):i180–i188, 2013.

Pier Francesco Palamara, Todd Lencz, Ariel Darvasi, and Itsik Pe’er. Length distributions of identity by descent reveal fine-scale demographic history. The American Journal of Human Genetics, 91(5):809–822, 2012.

Peter Ralph and Graham Coop. The geography of recent genetic ancestry across europe. PLoS Biology, 11(5):e1001555, 2013.

Harald Ringbauer, Graham Coop, and Nicholas H Barton. Inferring recent demography from isolation by distance of long shared sequence blocks. Genetics, 205(3):1335–1351, 2017.

Pauli Virtanen, Ralf Gommers, Travis E. Oliphant, Matt Haberland, Tyler Reddy, David Cour-napeau, Evgeni Burovski, Pearu Peterson, Warren Weckesser, Jonathan Bright, Stéfan J. van der Walt, Matthew Brett, Joshua Wilson, K. Jarrod Millman, Nikolay Mayorov, Andrew R. J. Nelson, Eric Jones, Robert Kern, Eric Larson, C J Carey, İlhan Polat, Yu Feng, Eric W. Moore, Jake VanderPlas, Denis Laxalde, Josef Perktold, Robert Cimrman, Ian Henriksen, E. A. Quin-tero, Charles R. Harris, Anne M. Archibald, Antônio H. Ribeiro, Fabian Pedregosa, Paul van Mulbregt, and SciPy 1.0 Contributors. SciPy 1.0: Fundamental Algorithms for Scientific Computing in Python. Nature Methods, 17:261–272, 2020. doi: 10.1038/s41592-019-0686-2.

Ciyou Zhu, Richard H Byrd, Peihuang Lu, and Jorge Nocedal. Algorithm 778: L-BFGS-B:Fortran subroutines for large-scale bound-constrained optimization. ACM Transactions on mathematical software (TOMS), 23(4):550–560, 1997.

